# Multiple domains of scaffold Tudor protein play non-redundant roles in *Drosophila* germline

**DOI:** 10.1101/2025.03.13.643173

**Authors:** Samuel J. Tindell, Alyssa G. Boeving, Julia Aebersold, Alexey L. Arkov

## Abstract

Scaffold proteins play crucial roles in subcellular organization and function. In many organisms, proteins with multiple Tudor domains are required for the assembly of membraneless RNA-protein organelles (germ granules) in germ cells. Tudor domains are protein-protein interaction modules which bind to methylated polypeptides. *Drosophila* Tudor protein contains eleven Tudor domains, which is the highest number known in a single protein. The role of each of these domains in germ cell formation has not been systematically tested and it is not clear if some domains are functionally redundant. Using CRISPR methodology, we generated mutations in several uncharacterized Tudor domains and showed that they all caused defects in germ cell formation. Mutations in individual domains affected Tudor protein differently causing reduction in protein levels, defects in subcellular localization and in the assembly of germ granules. Our data suggest that multiple domains of Tudor protein are all needed for efficient germ cell formation highlighting the rational for keeping many Tudor domains in protein scaffolds of biomolecular condensates in *Drosophila* and other organisms.

## Introduction

Scaffold proteins play major roles in different cells by bringing together their different partner proteins to initiate the effective response to cellular signals and induce and maintain formation of membraneless organelles (DiRusso CJ et al, 2022, Good MC et al, 2011). Scaffold proteins often contain several protein-protein interaction modules that allow them to bind to different partners during signal transduction or the assembly of biomolecular condensates. In particular, in germ cells of many animals, scaffold proteins containing multiple Tudor (Tud) domains play crucial roles in the assembly of RNA-protein membraneless organelles referred to as germ granules (Arkov AL & Ramos A, 2010, Gao M & Arkov AL, 2013, Pamula MC & Lehmann R, 2024, Ramat A et al, 2024, Voronina E et al, 2011). Tud domain is about 60 amino-acid β barrel structure which has a binding pocket lined with aromatic amino acids that associate with methylated lysines or arginines of partner proteins (Simcikova D et al, 2023). In particular, *Drosophila* Tud protein contains the highest number of Tud domains known in a single protein (eleven) (Vo HDL et al, 2019, Wahiduzzaman et al, 2024) and it has not been determined whether all these domains play a role in Tud protein structure and function during germ cell development or some domains are redundant or exclusively utilized in non-germline (somatic) cell types (Tindell SJ et al, 2020).

Using forward genetics maternal mutant screens and directed mutagenesis of the *tud* transgene, which expressed C-terminal fragment of Tud, previous studies characterized mutations in several Tud domains (Arkov AL et al, 2006, Boswell RE & Mahowald AP, 1985, Liu H et al, 2010). Virtually all these mutations caused defects in germ cell formation and decreased the levels of Tud protein or changed the morphology, decreased the size and the number of germ granules (polar granules) assembled in posterior cytoplasm of the egg (germ plasm). However, there was a set of the N-terminal Tud domains (domains 2-6), which have not been characterized with previous mutational approaches and it has not been clear whether these domains contribute to the structure and function of Tud scaffold in germ cell formation.

In this work, we asked whether, similarly to other previously studied Tud domains of Tud scaffold, majority of these poorly characterized N-terminal domains individually contribute to the structure and function of Tud protein in a single developmental process of primordial germ cell formation. To this end, using CRISPR/Cas9 methodology, we introduced deletions of each of the domains 2-5 in the native *tud* locus and tested whether these deletions cause defects in germ cell formation, Tud expression levels, Tud localization to the posterior pole of the early embryos and polar granule assembly.

Our data showed that each of these poorly characterized single domains contributes to germ cell formation by regulating Tud amounts in the germline, Tud transport and localization to the germ plasm and the size of polar granules. Overall, this work and previous research provide evidence for the importance of nearly all Tud domains of Tud scaffold for the protein structure and function and suggests that each of the multiple Tud domains of Tud protein is utilized during germline development in *Drosophila*. Surprisingly, there appears to be no redundancy of different Tud domains in Tud scaffold during its involvement in the formation of primordial germ cells in early *Drosophila* embryo.

## Results

### Tud domain 2-5 mutants show reduction in the number of primordial germ cells in *Drosophila* embryos

Previous work has characterized mutations in several Tud domains of Tud protein and helped to establish the importance of specific individual domains of this scaffold protein for primordial germ cell formation, binding to polar granule components, polar granule assembly and morphology. In particular, mutations in Tud domains 1 (Arkov AL et al, 2006) and 7-11 (Arkov AL et al, 2006, Boswell RE & Mahowald AP, 1985, Liu H et al, 2010) (Fig. S1) caused reduction in the number of primordial germ cells formed in posterior of early embryos or germ plasm localization of Tud domain binding protein Aubergine (Aub). In addition, germ plasm of Tud domain 1 (*tud*^A36^), 7 (*tud*^4^) and 10 (*tud*^B42^) mutants (Fig. S1) was examined with electron microscopy (EM) and the reduction in size, number or abnormal morphology of the polar granules were detected in these mutants (Arkov AL et al, 2006, Boswell RE & Mahowald AP, 1985).

In order to obtain the comprehensive understanding of how Tud protein uses its Tud domains during polar granule assembly and germ cell formation, using CRISPR methodology, we generated small deletions removing most of the remaining domains that have not been characterized previously (domains 2-5) in native *tud* locus (Fig. 1). While, similarly to Tud domains 2-5, Tud domain 6 was not well-characterized, we could not generate a similar deletion of this domain despite rigorous efforts.

**Figure 1.**
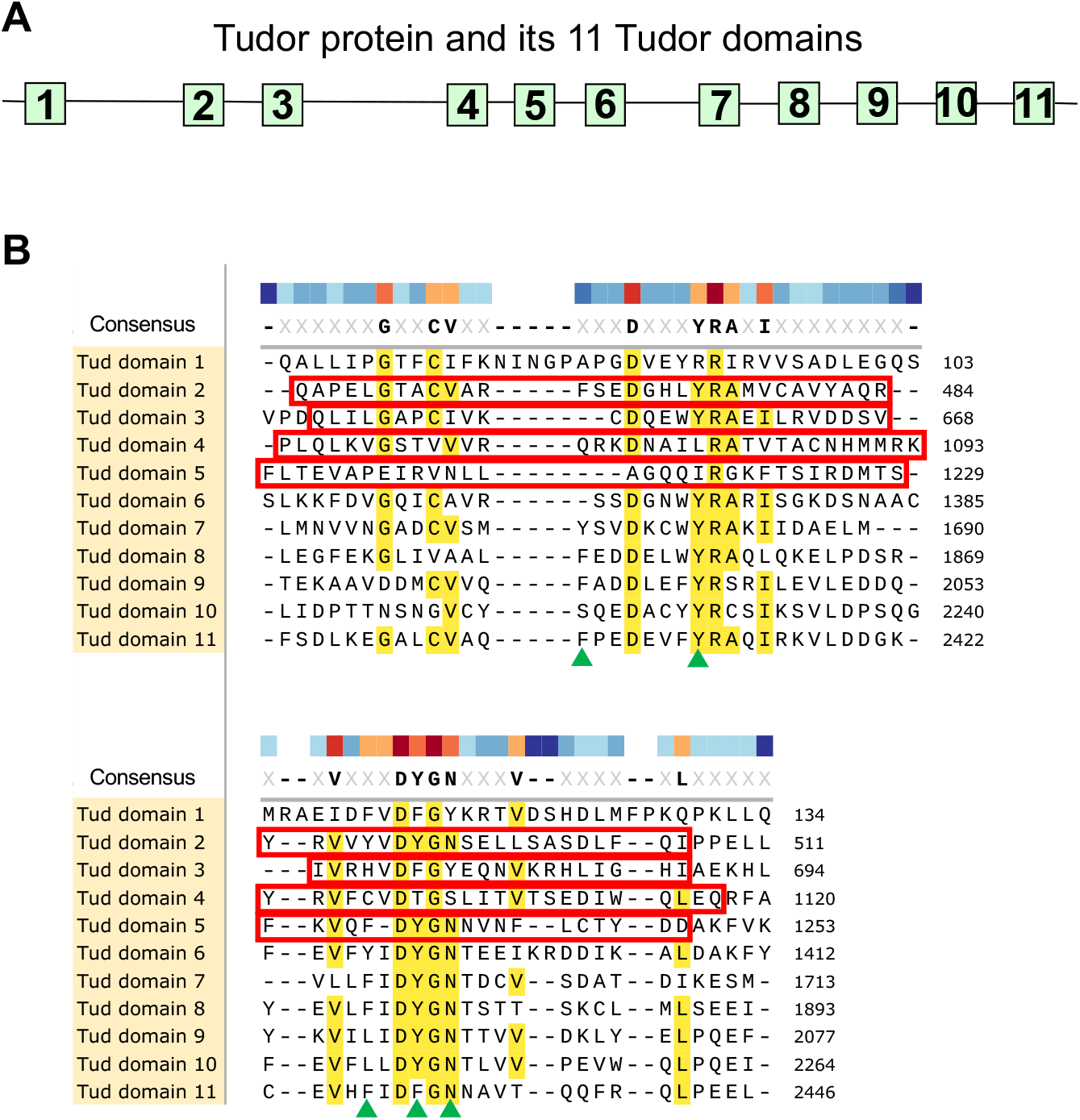
Tudor domains of Tudor scaffold protein and indicated deletion mutants in domains 2-5 generated by CRISPR methodology. **(A)** A diagram of Tud protein with its 11 Tud domains (squares) at their approximate locations is indicated. **(B)** Alignment of Tud domains of Tud protein. Conserved or similar residues are indicated in yellow and in consensus sequence. Poorly characterized Tud domains 2-5, analyzed in this study, were deleted by CRISPR methodology in the native *tud* locus and these deletions are outlined with red boxes. Green triangles point to the residues of the binding pocket of Tud domain 11 structure which binds to methylated arginine of Piwi protein Aubergine (Liu H et al, 2010).

First, we determined if each of the mutants shows reduction in the number of primordial germ cells formed in the embryos. Figure 2 shows that mutations in all of the domains 2-5 caused significant reduction in the number of germ cells. The strongest germ cell formation phenotypes were exhibited by Tud domain 3 mutant (*tud*^dom3^; no germ cells formed in 100% embryos examined) and Tud domain 4 mutant (*tud*^dom4^; only 12% embryos formed some germ cells). Tud domain 2 (*tud*^dom2^) and 5 (*tud*^dom5^) mutants showed less strong but still significant reduction in the number of primordial germ cells (about two-fold reduction in germ cell number compared to the wild-type control) (Fig. 2). These data indicate that presence of each domain is needed to maximize the involvement of Tud scaffold in germ cell formation.

**Figure 2.**
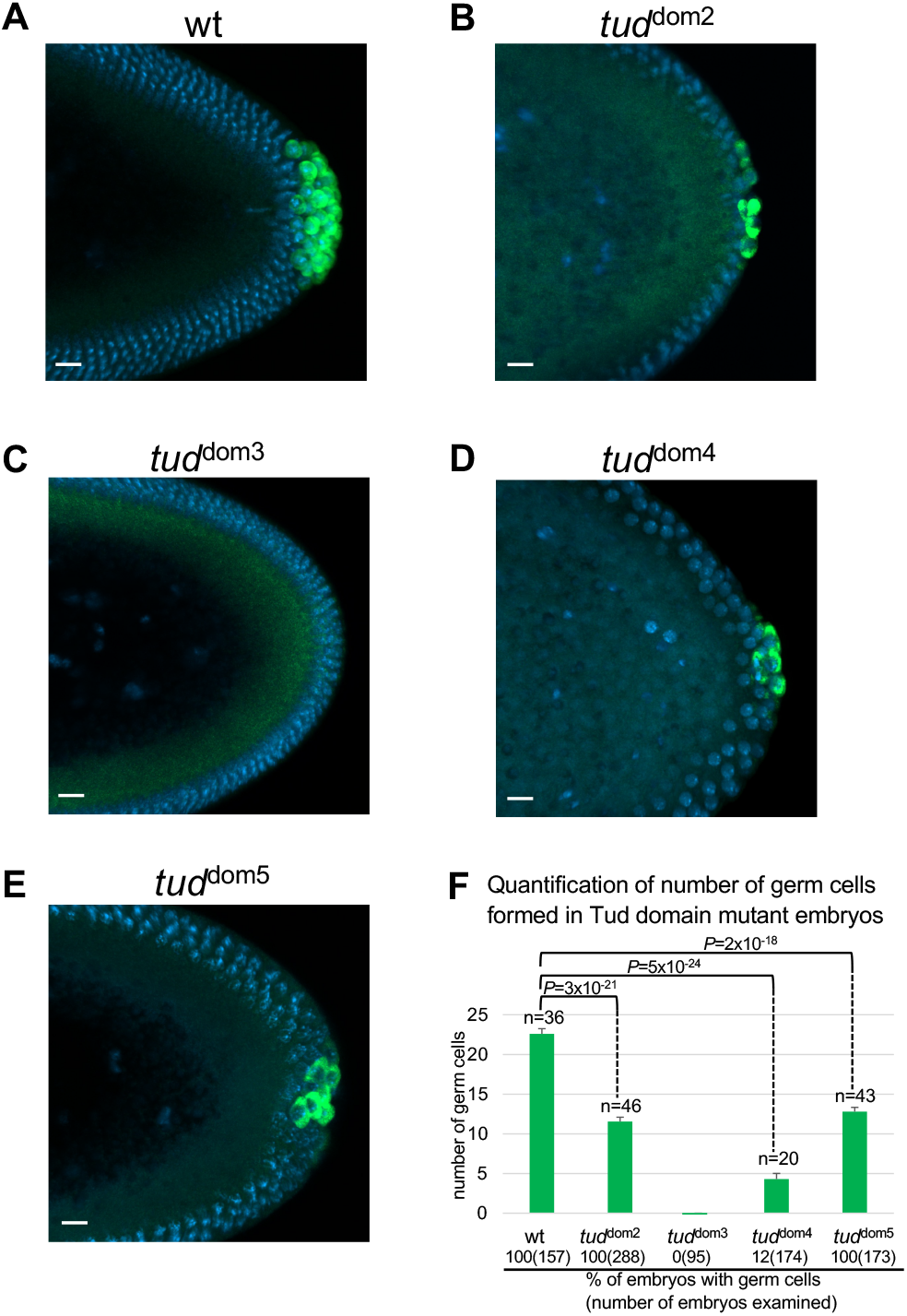
Mutations in Tudor domains 2-5 of Tudor scaffold protein cause defects in primordial germ cell formation in *Drosophila* embryos. **(A-E)** Germ cells formed at posterior pole of early blastoderm embryos are labeled with anti-Vasa (Vas) antibody (green). DAPI labels nuclei (blue). **(A)** Wild-type (*wt*) control embryos were produced by females with *wt tud* allele/*tud* deletion (*Df(2R)Pu*^*rP133*^, Materials and Methods). This *wt tud* allele was tagged with FLAG tag in the native *tud* locus with CRISPR editing (Tindell SJ et al, 2020) and was used for the production of all Tud domain mutants with CRISPR methodology reported in this work. **(B-E)** Mutant embryos were generated by females transheterozygous for the indicated Tud domain mutation and *tud* deletion *Df(2R)Pu*^*rP133*^. **(F)** Quantification of germ cells’ number (mean±s.e.m) formed in different Tud domain mutant embryos and *wt* control whose representative images are shown in **(A-E)**. Germ cells were counted from multiple embryos (*n*) at stages 10-13 of embryonic development. During these stages, the germ cells are spread out to a maximal degree inside the embryo and can be counted most accurately (Santos AC & Lehmann R, 2004). Germ cell numbers for *wt, tud*^dom2^, *tud*^dom4^ and *tud*^dom5^ mutants were 22.6±0.7, 11.5±0.6, 4.3±0.7 and 12.8±0.5 respectively. Reduction of germ cells number in all the mutants compared to *wt* control was statistically significant (unpaired two-tailed *t*-test was used; *P* values indicated). Separately, for each mutant, the percentage of embryos at stages 5-14 that contain any germ cells was scored and indicated at the bottom of the figure. While all *wt, tud*^dom2^ and *tud*^dom5^ embryos showed germ cells, only 12% of *tud*^dom4^ embryos formed germ cells and *tud*^dom3^ embryos failed to form any germ cells. Number of embryos scored is indicated in parentheses following the percentage value. In **(A-E)** scale bars are 10 μm.

### Mutations in Tudor domains affect Tudor protein enrichment in the germ plasm and protein expression levels

Defects in germ cell formation might be caused by the inability of Tud protein to reach the germ plasm or its reduced stability. Therefore, we determined whether Tud domain mutant proteins can be detected in the germ plasm of early preblastoderm embryos before germ cell formation. Figure 3 shows that Tud domain 2, 3 and 5 mutants fail to enrich Tud in germ plasm labeled with anti-Vasa (Vas) antibody (Fig. 3B, C, E). Contrary to this, Tud domain 4 mutant protein was able to specifically localize to the germ plasm (Fig. 3D) similar to the wild-type control (Fig. 3A).

**Figure 3.**
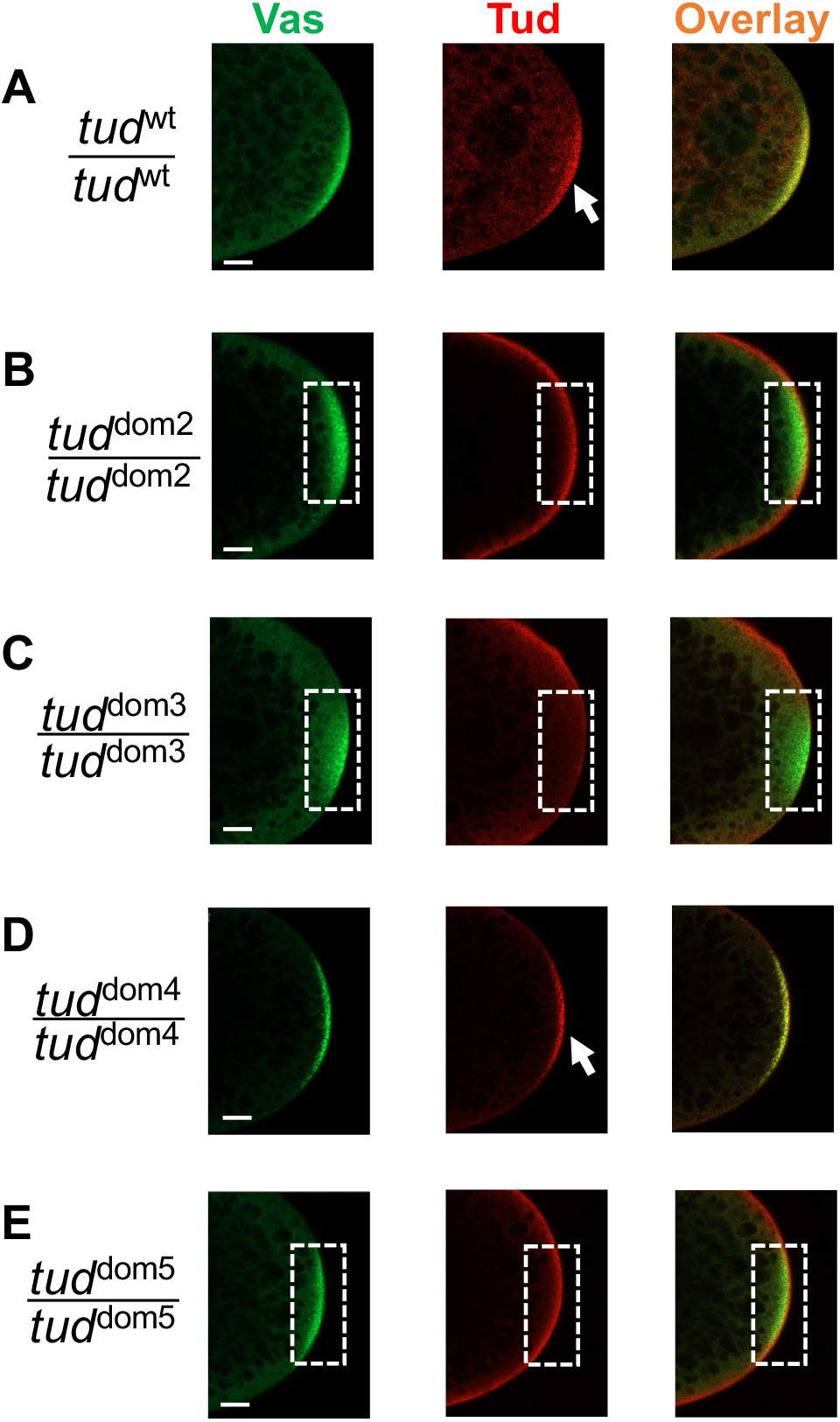
Different Tudor domain mutants show different levels of enrichment of Tudor protein in germ plasm of preblastoderm embryos. **(A-E)** Representative confocal microscopy optical sections of posterior poles of early preblastoderm embryos from homozygous wild-type (*wt*) *flag*-*tud* **(A)** or Tud domain mutant **(B-E)** females. The embryos were immunostained with antibodies against Vas protein (green) to label germ plasm and anti-FLAG antibody to label Tud (red). Overlay images are shown. While *wt* and Tud domain 4 mutant embryos show the localization of Tud protein to the germ plasm (arrows), Tud domain 2, 3 and 5 mutants fail to show Tud enrichment there (germ plasm area is outlined with dashed line) suggesting defects in transport to/maintenance in the germ plasm or in the expression of Tud protein. For each mutant and wild-type control experiments, z-stacks (18-49 optical sections in each stack) for each of the 9-10 preblastoderm embryos per genotype were acquired for analysis of protein distribution. Scale bars are 15 μm.

Next, we asked if *tud* mutants affected Tud protein levels. To this end, to determine whether the mutations decreased Tud stability or expression levels we used our new anti-Tud antibody (Fig. S2) and anti-FLAG antibody, since all CRISPR-edited *tud* alleles encoded N-terminal FLAG tag (Fig. 4A-C). While Tud protein amounts were reduced to about 40-60% of the wild-type control Tud levels in Tud domain 2, 4 and 5 mutants, that reduction cannot explain the lack of Tud accumulation in the germ plasm of Tud domain 2 and 5 proteins (Figs. 3B, E) since similarly expressed Tud domain 4 mutant protein (49% expression of the wild-type control) shows robust Tud enrichment in the germ plasm (Fig. 3D). Therefore, Tud domain 2 and 5 may contribute to the localization or maintenance of Tud in the posterior germ plasm.

**Figure 4.**
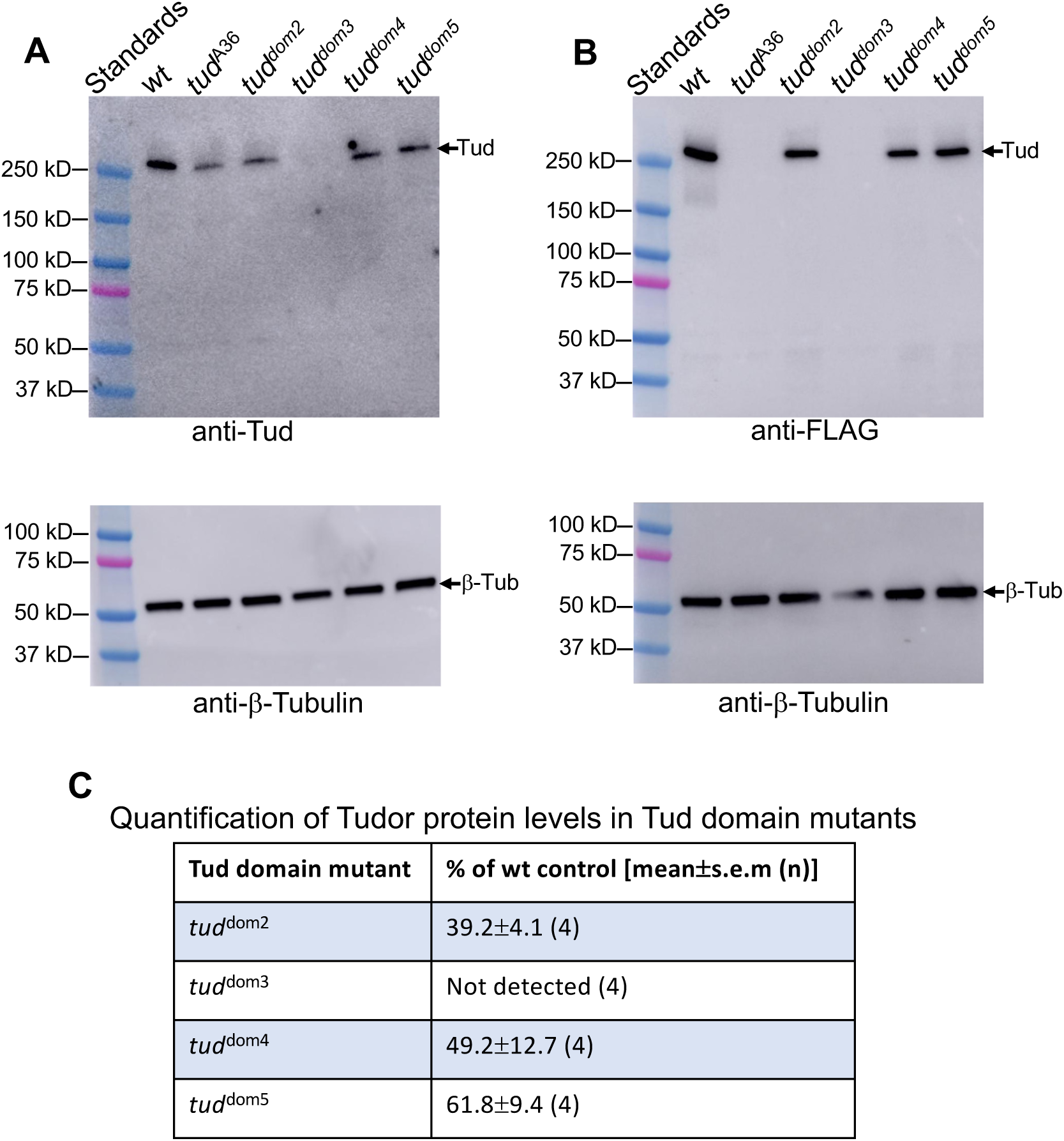
Expression levels of Tud domain mutants. **(A, B)** Western-blot data show expression levels of Tud protein in wild-type (*wt*) control and different Tud mutant ovaries isolated from females transheterozygous for a given *tud* mutant allele and *tud* deletion *Df(2R)Pu*^*rP133*^. In addition to Tud domain 2-5 mutant and the *wt tud* control genes, which all encode the N-terminal FLAG tag, previously characterized and quantified Tud domain 1 mutant (*tud*^A36^) (Arkov AL et al, 2006) (Fig. 1S, Table 1) was included in the western-blot experiments. This *tud*^A36^ does not have a FLAG tag. **(A)** Top: western-blot experiment with anti-Tud antibody (validated in Fig. S2) detects Tud expression in all mutants except for Tud domain 3 mutant. Bottom: Sample loading was controlled with anti-β-Tubulin antibody using the same gel shown in the top panel. **(B)** A different western-blot experiment with anti-FLAG antibody supports data obtained with anti-Tud antibody. Tagless *tud*^A36^ mutant protein serves as a negative control in this experiment. Top and bottom panels show western-blot data with anti-FLAG and anti-β-Tubulin antibody (loading control) respectively. **(C)** Expression levels of mutant Tud domain proteins were quantified as percentages of *tud* expression in *wt* controls (mean±s.e.m) from four biological replicate western-blot experiments using anti-Tud antibody.

Tud domain 3 mutant protein failed to be expressed at detectable levels (Fig. 4A-C), indicating that the mutation caused dramatic decrease in protein stability consistent with very strong germ cell formation phenotype for this mutant (Fig. 2C, F) and apparent lack of the mutant protein in the germ plasm (Fig. 3C). This is a striking result given that in this Tud domain mutant only one out of eleven Tud domains is deleted and it resembles previous data demonstrating a strong reduction in Tud protein amount when two aromatic amino acids in a single Tud domain 7 were mutated in the Tud fragment containing Tud domains 7-11 (Fig. S1) (Liu H et al, 2010).

**Table 1.**
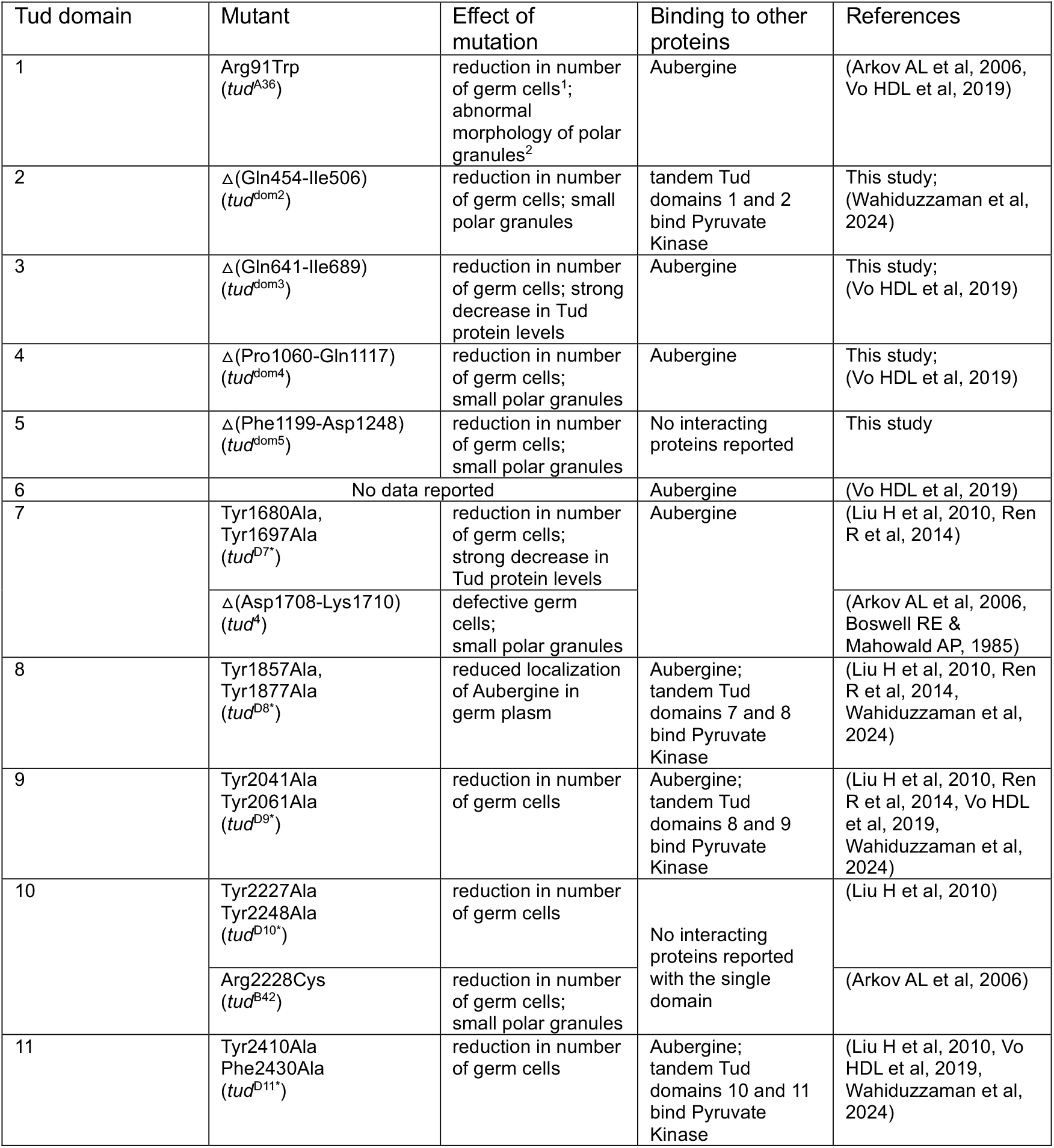
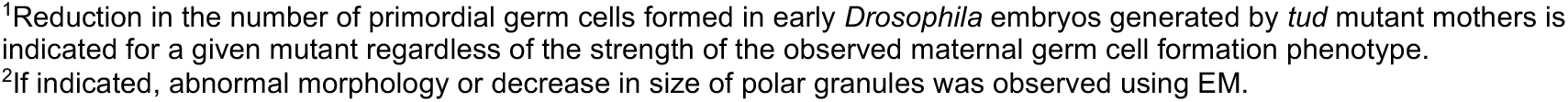
Role of Tudor domains in Tudor scaffold protein.

### Tud domain mutants affect polar granule assembly

The expression of Tud protein is essential for the assembly of polar granules, composed of RNA and proteins required for germ cell formation (Arkov AL et al, 2006, Boswell RE & Mahowald AP, 1985, Thomson T & Lasko P, 2004). Therefore, using super-resolution microscopy imaging and EM, we tested if Tud domains 2, 4, and 5 mutants, which all express Tud protein (Fig. 4) but are defective in germ cell formation (Fig. 2), show defects in polar granules size or morphology.

First, using super-resolution microscopy imaging, we detected incorporation of Tud and Vas proteins in small granules at the posterior poles of preblastoderm embryos in all these Tud domain mutants (Fig. 5A-D, three left panels).

**Figure 5.**
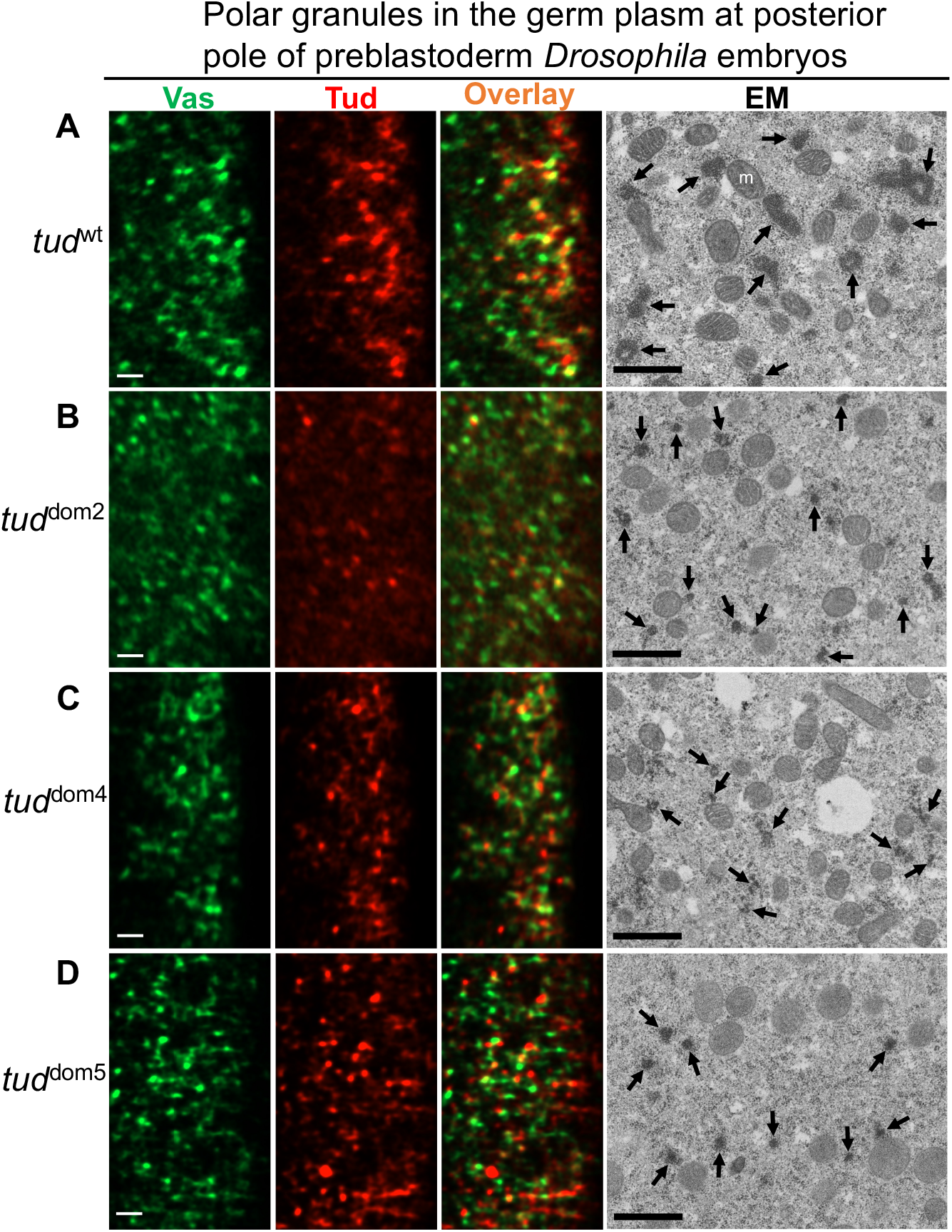
Tudor domain mutants assemble small electron-dense polar granules in germ plasm at posterior pole of early *Drosophila* embryos. **(A-D)** Germ plasm at the posterior pole of preblastoderm embryos was imaged with super-resolution microscopy (left three columns) and transmission electron microscopy (EM, right column) for *wt* control **(A)** and indicated Tud domain 2, 4, 5 mutants **(B-D)** shown to express Tud protein (Fig. 4). In these experiments, embryos were from females transheterozygous for an indicated *tud* mutant allele and *tud* deletion *Df(2R)Pu*^*rP133*^ and all Tud proteins had FLAG tag. For super-resolution experiments, embryos were immunostained with anti-Vas (green) and anti-FLAG (red) antibodies. Both Vas and Tud are assembled into granules in the germ plasm of the mutant embryos. Representative super-resolution optical sections for each mutant and *wt* control are shown. For super-resolution imaging, z-stacks for 8-10 embryos per genotype (21-101 optical sections per embryo) were analyzed. Consistent with super-resolution imaging, EM data further revealed electron-dense polar granules assembled in the mutants that were smaller than those in the *wt* control (indicated with arrows). Scale bars for all images are 1 μm, m, mitochondria.

Next, we determined if polar granules in the mutants can be visualized with EM approach. Specifically, in wild-type germ plasm polar granules were easily detected as characteristic amorphous membraneless electron-dense particles (Fig. 5A, right panel). Furthermore, Tud domain 2, 4, 5 mutants clearly assemble electron-dense polar granules, which, however, were smaller than in the wild-type control (Fig. 5B-D). Therefore, despite the apparent ability of Tud to be incorporated in the granules in germ plasm in all these Tud domain mutants, the mutations appear to cause reduction in size or electron density of polar granules.

## Discussion

In this work we provide evidence for non-redundant functions of multiple domains of Tud scaffold protein in primordial germ cell formation in *Drosophila*. Tud protein contains eleven domains and, based on previous research and this work, nearly all Tud domains of this protein are now shown to contribute to this specific developmental process during germline development (Table 1).

In particular, previous work showed the functional importance of Tud domains 1, 7-11 in different aspects of germ cell formation, interactions with methylated Tud protein binding partners Aub and Pyruvate Kinase (PyK) and the assembly of polar granules (Arkov AL et al, 2006, Gao M et al, 2015, Liu H et al, 2010, Vo HDL et al, 2019, Wahiduzzaman et al, 2024). However, apart from the contribution of some of the N-terminal Tud domains 2-6 in Aub binding detected in vitro (Creed TM et al, 2010, Vo HDL et al, 2019), the in vivo role of these domains in germ cell formation remained unknown, and it was not clear whether there is a functional redundancy among Tud domains of Tud scaffold in germ cell formation given their remarkably high number in a single protein. Also, since Tud protein was detected in soma (brain glia) (Tindell SJ et al, 2020), one could propose that there are different subsets of Tud domains each specific for Tud function in either soma or germline.

In this work, we generated small deletions in *tud* locus with CRISPR methodology that separately removed Tud domains 2-5, and characterized these novel mutants in detail using phenotypic analysis, super-resolution microscopy and EM. All these mutations caused defects in primordial germ cell formation. In addition, we demonstrated that these Tud domain mutants had different effects on Tud expression and enrichment in the posterior germ plasm. In particular, Tud domain 2 and 5 mutant preblastoderm embryos showed no Tud enrichment in the posterior pole, and deletion of Tud domain 3 resulted in no detectable Tud protein expression explaining complete absence of germ cells in this mutant. In contrast to Tud domain 2 and 5 mutants, Tud domain 4 mutant showed strong enrichment in the germ plasm. However, despite this germ plasm enrichment and contrary to Tud domains 2 and 5 mutants, Tud domain 4 mutation caused a very strong reduction in the number of germ cells. The difference in the strength of the germ cell formation phenotype among Tud domain 4 mutant and the other mutants cannot be explained by the different expression levels of Tud since Tud domain 2-, 4- and 5-mutant proteins are expressed at similar amounts. Our data suggest distinct function of the N-terminal Tud domains: Tud domains 2 and 5 might be similarly involved in Tud protein transport or enrichment in the germ plasm and Tud domain 4 is needed downstream of Tud localization step leading to germ cell formation.

In all *tud* mutants, that were examined for the presence of polar granules, these membraneless organelles are either extremely difficult to find or smaller or of abnormal morphology than in the wild-type control (Arkov AL et al, 2006, Boswell RE & Mahowald AP, 1985, Thomson T & Lasko P, 2004). In this work, using EM approach to visualize polar granules, we found that all Tud protein-expressing Tud domains 2, 4 and 5 mutants formed small electron-dense polar granules, which appear to contain mutant Tud protein as detected with super-resolution microscopy imaging. The size reduction of polar granules in the mutants may contribute to the reduction in germ cell number detected in these mutants as the granule components are essential for germline development. Since single Tud domains and their methylated ligands can drive the formation of biomolecular condensates (Courchaine EM et al, 2021), it is conceivable that the lack of certain single Tud domains in Tud scaffold may hinder the formation of polar granules. However, it is remarkable that a single domain has a noticeable effect on polar granule assembly given that ten remaining Tud domains are still present in Tud scaffold.

Future research will provide biochemical and structural understanding of how all Tud domains and their specific binding partners work together on Tud scaffold as one functional system and shed light on molecular rational for the non-redundancy of multiple Tud domains indicated by this work.

## Materials and Methods

### Tudor domain mutants

Deletions of Tud domains 2-5 were generated in the native *tud* locus that was previously tagged with N-terminal FLAG tag (Tindell SJ et al, 2020) using CRIPR/Cas9 methodology. The Tud domain deletions and the corresponding *tud* alleles are shown in Fig. S1 and are as follows. *tud*^dom2^: Gln454-Ile506; *tud*^dom3^: Gln641-Ile689; *tud*^dom4^: Pro1060-Gln1117; and *tud*^dom5^: Phe1199-Asp1248. All deletions were generated using oligonucleotide (ODN) donor templates for CRISPR/Cas9-mediated homology-directed repair (HDR) as previously described (Gratz SJ et al, 2015). Mutant lines were generated by Rainbow Transgenic Flies, Inc. (Camarillo, CA, USA) and confirmed with sequencing. Unless specified otherwise, for most experiments described in this work, embryos or ovaries were from females transheterozygous for a *tud* allele and a *tud* deletion, *Df(2R)Pu*^*rP133*^.

### Production of rabbit anti-Tud antibody

Rabbit anti-Tud antibody was raised against purified C-terminal Tud protein fragment containing Tud domains 7-11 (amino acids 1605-2515). This Tud fragment was produced from pET SUMO vector in *E. coli* BL21 (DE3) cells as described previously (Wahiduzzaman et al, 2024). The antibody was produced by Cocalico Biologicals (Denver, PA, USA) according to the standard protocol detailed previously (Kharel K et al, 2024). Subsequently, the antibody was validated and confirmed to recognize Tud protein specifically in both western-blot (Fig. 4) and immunohistochemistry (Fig. S2) experiments. For immunohistochemistry, 1:1500 dilution of the antibody was used.

### Immunohistochemistry

These methods have been described (Navarro C et al, 2004, Stein JA et al, 2002). For whole-mount immunostaining of fly embryos, rabbit anti-Vasa antibody (1:1000) (Stein JA et al, 2002) was used. Also, in order to detect Tud, mouse anti-FLAG antibody (Millipore Sigma, 1:2500) was used.

### Analysis of mutant Tud protein expression

Expression levels of Tud protein in ovarian extracts from *tud* mutants and wild-type control, were quantified with western-blot procedure as detailed previously (Arkov AL et al, 2006). The following antibodies were used for detection of Tud protein: rabbit anti-Tud validated in this work (1:1500), mouse anti-FLAG antibody (Millipore Sigma, 1:3000) as alternative antibody to confirm Tud expression, and mouse anti-β-Tubulin antibody (Millipore Sigma, 1:5000) as a loading control.

### Super-resolution microscopy

Super-resolution microscopy imaging was carried out essentially as described (Wahiduzzaman et al, 2024). In particular, a Zeiss LSM 980/Airyscan super-resolution module system, inverted laser scanning confocal microscope AxioObserver and Plan-Apochromat x63/1.4 Oil DIC M27 objective were used. For every experiment, images were acquired equally for mutants and wild-type control as z-stacks, and subsequently analyzed with Imaris software (version 9.5, Oxford Instruments) and an HP Z8 workstation.

### Electron microscopy

Preparation of *Drosophila* embryos to image posterior germ plasm and polar granules with EM was done essentially as described (Arkov AL et al, 2006). In particular, after initial fixation of the dechorionated embryos in heptane saturated with 12.5% glutaraldehyde for 20 minutes at room temperature, vitelline membranes were removed manually. Then, the embryos were fixed in 2% paraformaldehyde/2.5% glutaraldehyde in 0.1M Sodium Cacodylate Buffer (pH 7.4) overnight at 5°C. The fixed embryos were rinsed three times for 10 minutes with 0.1M Sodium Cacodylate Buffer followed by staining with 1% Osmium Tetroxide for 1 hour. Samples were rinsed one time for 10 minutes in 0.1M Sodium Cacodylate Buffer and stained en bloc with 1% Uranyl Acetate for 30 minutes. Samples were rinsed twice with distilled water then dehydrated progressively with 30%, 50%, 70%, 90% and 95% ethanol for 10 minutes and three times with anhydrous ethanol for 15 minutes. Embedding was performed with 1:1 Spurr’s Resin and anhydrous ethanol along with an increased concentration of 3:1 resin and anhydrous ethanol for one hour. Samples were placed in 100% resin under vacuum for 24 hours without agitation then cured at 70°C for 24 hours. The embedded samples were sectioned to 80 nm with a Leica UC7 ultramicrotome and placed onto nickel slot grids with formvar support film. Post lead citrate staining was performed for 2 minutes on the grids and followed by rinsing with DI water. The samples were imaged with a Hitachi HT-7700 TEM at 80 kV.

## Acknowledgements

We thank Wahiduzzaman for the production of Tud domains 7-11 fragment used for the generation of anti-Tud antibody and E. Hackney for his comments on the manuscript. This study was supported by a grant from National Science Foundation MCB-2130162 to ALA. Also, this work was supported in part by a grant from the NIH National Institute of General Medical Sciences, P20GM103436.

## Author Contributions

SJ Tindell: methodology, investigation, data analysis, manuscript review.

AG Boeving: investigation, data analysis, manuscript review.

J Aebersold: methodology, investigation, data analysis, manuscript review.

AL Arkov: conceptualization, methodology, investigation, data analysis, funding acquisition, writing – original draft.

