## Supplementary information for "Multiple domains of scaffold Tudor protein play non-redundant roles in *Drosophila* germline"

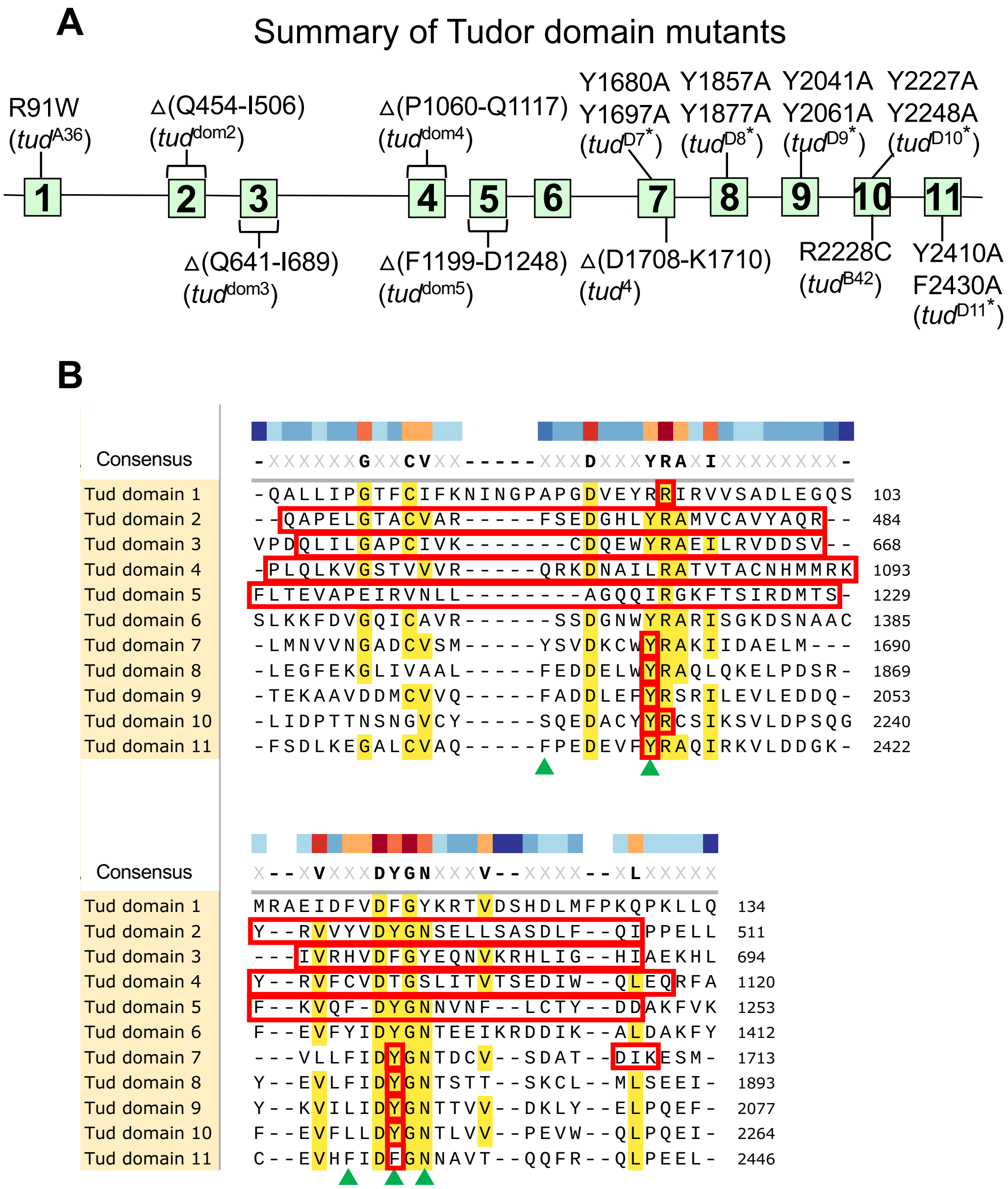


**Figure S1. Summary of mutations in Tudor domains of Tudor protein.**
**(A)** A diagram of Tud protein with all known Tud domains' mutations indicated in domains 1 (Arkov AL et al, 2006), 2-5 (this study) and domains 7-11 (Arkov AL et al, 2006, Boswell RE & Mahowald AP, 1985, Liu H et al, 2010). Deletion mutations are indicated as '∆'. Contrary to domains 2-5, CRISPR approach used in this study, failed to generate a comparable deletion of Tud domain 6 and also, previous mutational approaches did not isolate a mutation in this Tud domain. **(B)** Alignment of Tud domains shows the residues mutated in all Tud domain mutants shown in **(A)** (indicated in red). Part of this Figure, which shows the alignment, deletion mutations of Tud domains 2-5 and residues of methylarginine-binding pocket of Tud domain 11 (green triangles) is shown in Fig. 1B.

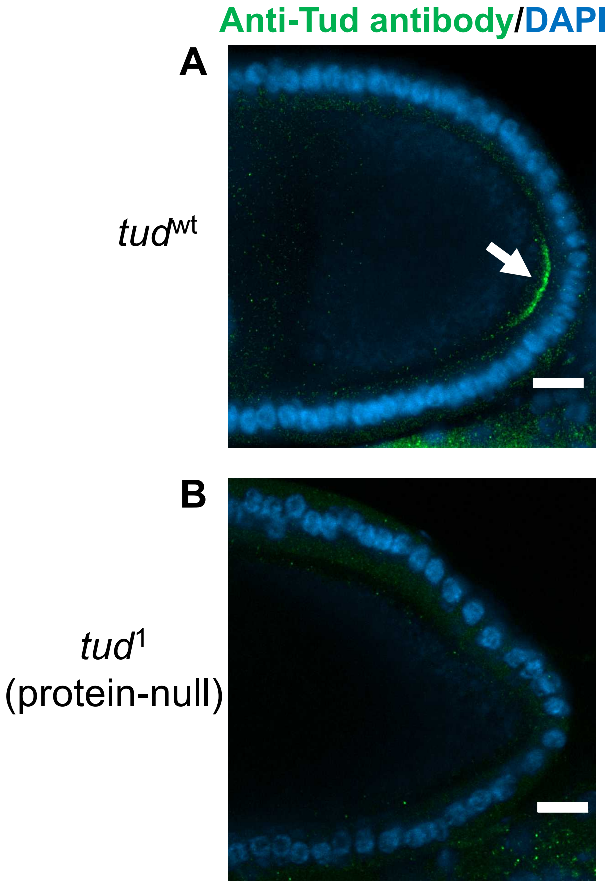


**Figure S2. Validation of rabbit anti-Tudor antibody.**
**(A)** Tud protein in the germ plasm at the posterior pole of the wild-type oocyte (*tud*$^{wt}$) at a mid-oogenesis stage is labeled with new rabbit anti-Tud antibody (green, arrow, Materials and Methods). Nuclei of follicle cells surrounding the oocyte are labeled with DAPI. For these immunostaining experiments, ovaries from *tud*$^{1}$/*CyO*, which carry one copy of *wt tud* allele on *CyO* balancer chromosome along with the original protein-null *tud*$^{1}$ mutant allele (E. Wieschaus and C. Nüsslein-Volhard, unpublished; sequenced in (Arkov AL et al, 2006)), were used. **(B)** Specificity of the antibody was further confirmed in immunostaining experiments using ovaries from females transheterozygous for *tud*$^{1}$ and *tud* deletion *Df(2R)Pu*$^{rP133}$. As expected, no Tud was detected in the germ plasm of the oocytes in the *tud*$^{1}$ mutant ovaries. In addition, this antibody was validated in western-blot experiments (Fig. 4). Scale bars are 15 μm.
